# Working memory shapes neural geometry in human EEG over learning

**DOI:** 10.1101/2025.01.21.634110

**Authors:** Michał J. Wójcik, Amy Li, Dante Wasmuht, Jake P. Stroud, Mark G. Stokes, Nicholas E. Myers, Laurence T. Hunt

**Affiliations:** Department of Experimental Psychology, University of Oxford, Oxford, UK; Department of Physiology, Anatomy and Genetics, University of Oxford, Oxford, UK; Department of Engineering, University of Cambridge, Cambridge, UK; School of Psychology, University of Nottingham, Nottingham, UK; Department of Psychiatry, University of Oxford, Oxford, UK

## Abstract

Working memory has been traditionally studied as a passive storage for information. However, recent advances have suggested that working memory is prospective rather than retrospective, meaning that its content undergoes transformations that will support future behaviour. One perspective that underscores this notion conceptualises memory processes as a computational resource that can be used to reduce the complexity of computation at decision time. Here, we explore this perspective by examining whether the process of maintenance shapes neural geometry and leads to low-dimensional representations during storage and later decision time. We recorded EEG in 25 human participants who learnt to solve a XOR task. We hypothesised that separating task features by a working memory delay would result in participants temporally decomposing the XOR computation, by prospectively processing one of the task features early in trial time. In line with our predictions, participants transformed the first feature from a sensory to an abstract format and maintained this pre-processed information throughout the delay. This process was related to the low-dimensional representation required at decision time early in learning, a representation that has recently been shown to support later cross-generalisation. These results demonstrate that low-dimensional representations, elsewhere associated with slow learning, might also provide a mechanism for maintenance processes in working memory.

## Introduction

Working memory, the capacity to briefly retain and manipulate information, is a foundational element of adaptive and flexible behaviour. A dominant view has emphasised the role of working memory in storing past sensory input (Goldman-Rakic, 1995). Yet there is also a clear role for working memory being used in anticipation of future task demands, supported by both empirical (Quintana & Fuster, 1992; Rainer et al., 1999; Sakai & Miyashita, 1991; Spaak et al., 2017; Stokes, 2015; Stokes et al., 2013; Wolff et al., 2017) and theoretical studies (Stroud et al., 2023, 2024). In particular, electrophysiological research in humans and primates has revealed that information in working memory undergoes transformations, yielding abstract representations that guide future behaviour, rather than merely mimicking sensory codes (Meyers et al., 2008; Spaak et al., 2017; Stokes, 2015; Stokes et al., 2013). The shift from sensory to abstract coding is now recognised as an inherent computational characteristic of working memory, possibly reflecting optimal loading of information into a working memory store (Stroud et al., 2023).

Insights from machine learning and computational neuroscience further highlight the idea that memory processes can be viewed as a resource for computations rather than a passive mechanism for storage (Dasgupta & Gershman, 2021; Ehrlich & Murray, 2022). In this light, working memory adapts computations to the current task demands (Dasgupta & Gershman, 2021); pre-computed information can be stored in working memory, and thus reduce the computation time at the moment of the decision (Braver, 2012; Hunt et al., 2021). This perspective is further supported by computational modelling of neural circuits that contends that working memory will change neural geometry in a way that supports the temporal decomposition of computations (Ehrlich & Murray, 2022). This work suggests that the computational load at the moment of action can be thus alleviated by decomposing complex operations into several simple problems solved sequentially in time.

One of the mechanisms of reducing neural dimensionality and thus the complexity of neural computations is filtering out irrelevant information through working memory selection (Jacob & Nieder, 2014). Extensive empirical and theoretical investigations demonstrate that low-dimensional representations are characterised by enhanced efficiency on various metrics (Barak et al., 2013; Bernardi et al., 2020; Cromer et al., 2010; Cueva et al., 2020; Flesch et al., 2022). For example, these representations can reduce the impact of noise on ongoing decision processes and provide robust and stable readouts (Fusi et al., 2016; Wójcik et al., 2023). The compressed format permits easy generalisation and thus increases adaptability (Chien et al., 2023; Reinert et al., 2021; Wójcik et al., 2023).

Using non-human primates, we recently demonstrated that low-dimensional representations are acquired over learning spanning multiple days (Wójcik et al., 2023). We found that metabolically constrained optimisation provides a parsimonious description of this process (Stroud et al., n.d.). Yet these observations were made under conditions where all necessary inputs were still available from the environment at the moment of the decision, rather than being sequentially presented to the participant and requiring retention and/or prospective manipulation over a delay. In an alternative setting, involving delay periods, low-dimensional representations could be constructed via working memory selection and prospective re-coding of features that enter working memory.

Here, we explored whether engaging working memory leads to the construction of low-dimensional neural representations. To test these predictions, we asked human participants to learn an exclusive-or (XOR) mapping – a problem that can be solved by a range of representations, from low- to high-dimensional (Ehrlich & Murray, 2022). Crucially, we presented the elements of the XOR mapping separately, with a temporal delay, to test what information was stored and transformed in working memory. By recording the EEG signal from naïve to expert performance on the task, we also tracked how the dimensionality and geometry of neural representations changed with learning. Based on theoretical work that suggests that working memory can be used to temporally decompose computations (Dasgupta & Gershman, 2021; Ehrlich & Murray, n.d.), we hypothesised that separating task features with a memory delay would result in the preparation of an abstract, low-dimensional context signal (representing the first task feature) before all necessary information for solving the task was presented. This decomposition should lead to a low-dimensional representation at the moment of the decision and be accompanied by tell-tale reaction time and performance biases (i.e., context switch costs). Specifically, we hypothesised that the task feature presented first would be used to contextualise the processing of subsequent features, and that trial-to-trial changes to this context would impact behavioural performance more than changes to subsequently presented features (Mayr & Kliegl, 2003; Vandierendonck et al., 2010).

Consistent with these predictions, participants rapidly learned to construct a contextual signal from sensory information provided by the first feature and maintained it throughout the delay period. Irrelevant dimensions of the contextual cue were suppressed early in the learning process. Notably, participants who exhibited stronger abstract coding of the first feature early in learning generated lower-dimensional representations at decision time. These low-dimensional representations closely resembled the low-dimensional representations observed in non-human primates after extensive training spanning several weeks (Wójcik et al., 2023). Interestingly, neural dimensionality did not decrease as a function of learning but remained low throughout, with reductions observed during trials associated with XOR coding failures. These findings suggest that neural dimensionality was not adapted as a learning strategy to achieve low-dimensional representations at decision time. Overall, this study highlights that low-dimensional representations, critical for complex decision-making, can be rapidly constructed via working memory selection processes.

## Results

Participants learned a cued stimulus-response mapping task with delay between context cue and stimulus-response mapping (**Fig. 1a, b**). First, a coloured circle was presented (colour). After a delay (fixation point), a shape was displayed, and participants were instructed to choose one of two response buttons (left or right). A nonlinear combination of the colour and shape predicted whether the left or the right response button was correct (the XOR rule). Importantly, to test whether irrelevant dimensions were discarded via working memory selection, the task consisted of two instances of the XOR rule (**Fig. 1c-d**). For example, in colour pair 1 (blue and green) the combination “blue-square” and combination “green-diamond” predicted the left response button to be correct. “Green-square” and “blue-diamond” predicted the right response button to be the correct one. In colour pair 2, the colours pink and khaki shared the same shape-response mapping as the colours blue and green in colour pair 1, respectively (e.g., “pink-square” and “khaki-diamond” were mapped onto the left response button). This allowed us to distinguish context coding (relevant dimension) from colour coding (irrelevant dimension) in the neural analyses.

**Figure 1.**
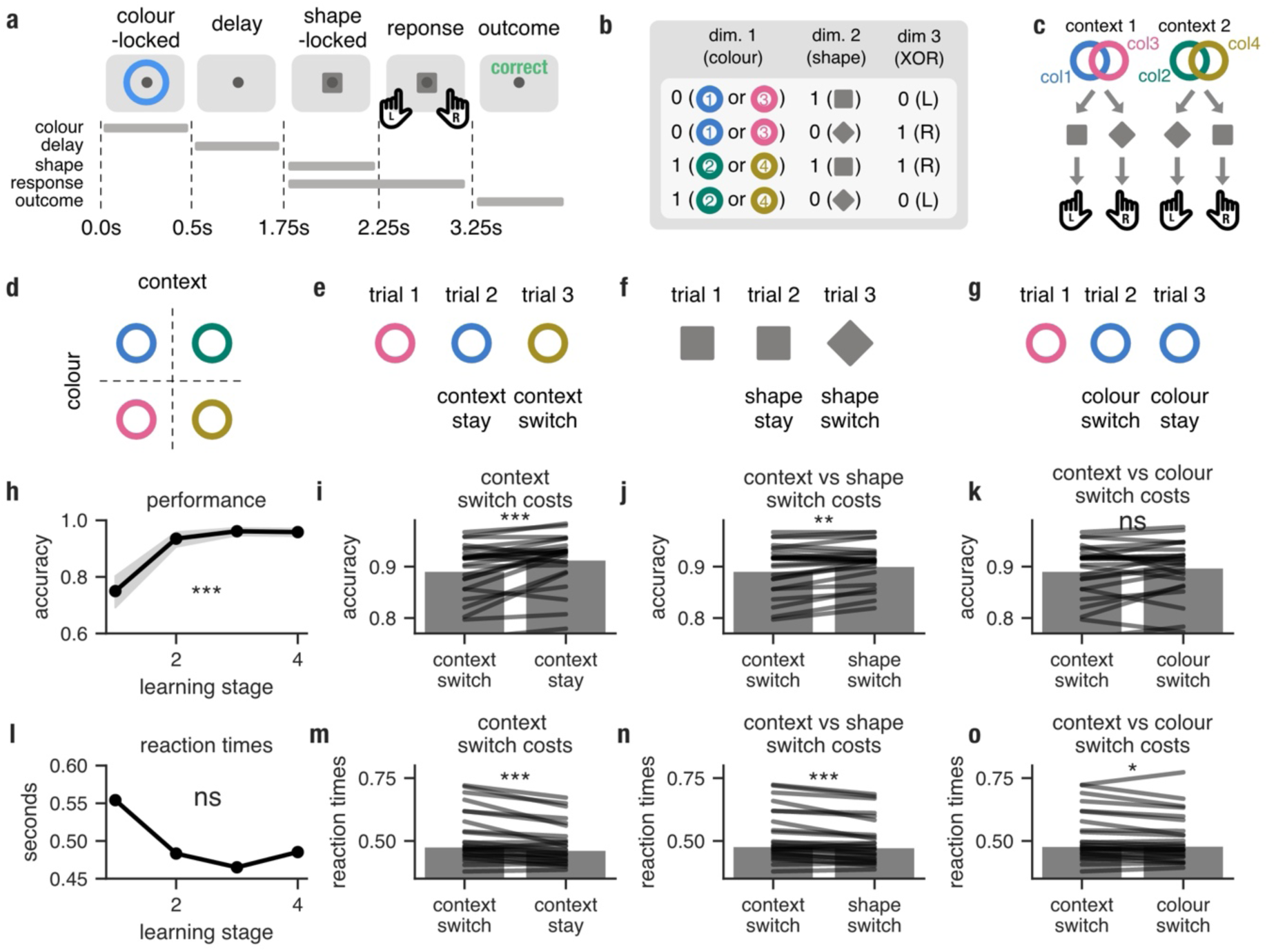
Experimental design and behavioural results. **a**, Timeline of task events in a single trial. **b**, Participants learnt by trial-and-error to combine two task features (colour and shape) in a non-linear fashion (XOR) and respond either by pressing the left or the right response button. For example, blue+square and green+diamond combined with left button press (XOR == False (0)) were followed by positive feedback; blue+diamond and green+square combined with right button press (XOR == True (1)) were also followed by positive feedback. Any other combination was followed by negative feedback. **b**, The task consisted of three features: colour (*n* = 4), shape (*n* = 2) and XOR conditions (*n* = 2; aligned fully with left and right button responses). **c**, Participants could generalise the meaning of colour 1 (e.g. blue) to colour 3 (e.g. pink), and vice versa, as both shared the same shape-response mapping. This could result in colours being grouped by the contextual information they provided for the shape-response mapping. **d**, The cue stimulus (first feature) had two dimensions: the task-irrelevant colour dimension and the task-relevant context dimension. **e**, In context switch trials the context of the current trial (trial 3) changed compared to the previous trial (trial 2) whereas in context stay trials the context of the current trial (trial 2) remained the same compared to the previous trial (trial 1). **f**, In shape switch trials the shape of the current trial (trial 3) changed compared to the previous trial (trial 2) whereas in shape stay trials the shape of the current trial (trial 2) remained the same compared to the previous trial (trial 1). **g**, In colour switch trials the colour of the current trial (trial 2) changed compared to the previous trial (trial 1) whereas in colour stay trials the colour of the current trial (trial 3) remained the same compared to the previous trial (trial 2). **h**, Mean performance accuracy plotted as a function of learning stages; shaded area indicates 95% CIs obtained through random resampling with replacement (*n* = 1000); accuracy in stage 1 was compared to accuracy in stage 4. **i**, Accuracy is lower on context switch trials (the context of the current trial changed compared to the previous trial) than context stay trials (the context of the current trial remained the same compared to the previous trial); horizontal lines are individual participants. **j**, Accuracy is also lower on context switch trials than shape switch trials (the shape of the current trial changed compared to the previous trial). Plotting conventions analogous to e. **k**, Comparison of context switch trials with colour switch trials (the colour of the current trial changed compared to the previous trial; as the context remained the same this measure captures changes caused by sensory differences only). **l-o**, Analogous to panels d-g but plotted for reaction time medians (before transformations). All p-values were calculated using t-tests for dependent samples (***, p < 0.01; **, p <0.01; *, p < 0.05; †, p < 0.1; n.s., not significant); reaction times were transformed using the log transformation prior to running parametric tests.

Participants needed to infer the correct responses through trial and error. As the task consisted of eight different stimuli-response combinations which formulated two instances of the same XOR rule, participants could have used two different strategies: (1) memorise all combinations (a high-dimensional strategy) or (2) capitalise on the colour feature being presented first and generate a low-dimensional representation at decision time, i.e., transform the colour information into an abstract context signal (context-sensitive strategy, **Fig. 1c**) before the shape onset to reduce the computational complexity at the moment of the decision. These strategies can be distinguished by their impact on trial-to-trial variability in reaction times and performance (Mayr & Kliegl, 2003; Vandierendonck et al., 2010). More precisely, if a strategy based on context information was used, the first feature would be considered more important and trial-to-trial changes in context (**Fig. 1e**) would have a stronger negative impact on behaviour (switch cost) than trial-to-trial changes in the second presented feature (shape; **Fig. 1f**). Conversely, with a memory-based strategy, both context and shape are equally important, leading to comparable switch costs for both context and shape switches. Our design further allowed us to distinguish costs associated with a context switch from costs arising from a mere change in the colour of the cue (**Fig. 1g**).

### Human behaviour exhibiting evidence of employing a context-sensitive strategy

Participants learned to solve the task rapidly with a performance above chance level (50%) in the first learning stage (i.e., first 25% of trials), *M* = .75, *t*(24) = 9.16, *p* < .001 (**Fig. 1h)**. Task accuracy also significantly increased with learning from stage 1 (*M* = .75) to stage 4 (*M* = .96), where participants exhibited ceiling-level performance, *t*(24) = 7.22, *p* < .001, *d* = 1.51. The learning-induced increase in performance was not accompanied by a significant decrease in reaction times, *M* = 554 *ms vs M* = 485 *ms*, *t*(24) = 1.15, *p* = 0.13, *d* = 0.19 (**Fig. 1l**).

We explored whether participants utilised a context-sensitive representation to solve the task by comparing context switch costs to shape switch costs. Performance in context switch trials (*M* = 89.2% ; where context in the current trial changed in comparison to context in the previous trial) was lower compared to context stay trials (*M* = 91.3%; where the same context was presented in the current and previous trial), *t*(24) = −3.85, *p* < .001, *d* = 0.3 (**Fig. 1i**). Context switches led to lower accuracy than shape switches, *M* = 89.2% *vs M* = 90.1%, *t*(24) = −2.96, *p* < .001, *d* = 0.13 (**Fig. 1j**). We found no differences in performance when context-switch trials and colour-switch trials were compared (when the previous colour was different from the current colour, but the context stayed the same) *M* = 89.2% *vs M* = 89.8%, *t*(24) = −1.18, *p* = 0.13, *d* = 0.08 (**Fig. 1k**), indicating that context switch performance costs may in part arise from cue switches.

Reaction time switch costs mirrored the pattern of results found in performance switch costs. Context switch trials were characterised by slower median reaction times compared to context stay trials, *M* = 478 *ms vs M* = 464 *ms*, *t*(24) = 4.34, *p* < .002, *d* = 0.22 (**Fig. 1m**) and shape switch trials, *M* = 479 *ms vs M* = 475 *ms*, *t*(24) = 4.42, *p* = 0.0, *d* = 0.15 (**Fig. 1n**). Interestingly, we also found that participants exhibited slightly slower reaction times in context switch trials as compared to colour switch trials, *M* = 480 *ms vs M* = 481 *ms*, *t*(24) = 2.07, *p* = 0.02, *d* = 0.07 (**Fig. 1o**).

These results indicate that participants used a highly structured representation of the task, in which the context was used to facilitate the subsequent processing of the shape feature. This led to worse and slower adaptation on trials where a switch in this higher-order feature took place. Next, we explored whether these behavioural effects translated into the hypothesised changes in neural codes.

### Humans learn to maintain the context information and construct the XOR feature over learning

To solve the task, participants were required to construct an XOR dimension by mixing nonlinearly the context and shape features. The initial cue stimulus, consisting of a coloured circle, could be classified along two dimensions: the irrelevant *colour* dimension, and the relevant *context* dimension (**Fig. 1d**). The former is retrospective in that it represents sensory differences between colours (blue vs pink or green vs khaki; **Fig. 1c**), whereas the latter is prospective: it dictates the context within which the next feature (shape) should be processed (‘square-left/diamond-right or square-right/diamond-left’; **Fig. 1c**). As the stimulus is displayed only for 500ms at the beginning of the trial, participants needed to maintain a representation in at least one of these formats in working memory until the shape was displayed. We hypothesised that across learning, participants would exhibit an increase in context coding but not colour coding in the delay period and that this would lead to an increase in XOR coding in the shape-locked period.

We first employed linear discriminant analysis (LDA) decoding on each time-point of the EEG signal separately, on trials pooled across learning stages. Context information decoding peaked early, in the colour-locked period, was maintained throughout the delay to then increase slightly after shape presentation, *cluster* 1: 0.082 − 1.188*s*, *p* = 0.001 and *cluster* 2: 1.208 − 3.2*s*, *p* = 0.001 (**Fig. 2a**). Next, the EEG signal was averaged in the colour-locked, delay-locked and shape-locked periods to explore learning-induced changes in decoding scores (see *Methods, decoding* for more details). We found that context coding in stage 4 increased as compared to stage 1 in the delay-locked period: *M* = 0.519 *vs M* = 0.507, *t*(24) = −1.8, *p* = 0.04, *d* = 0.35 (**Fig. 2b**).

**Figure 2.**
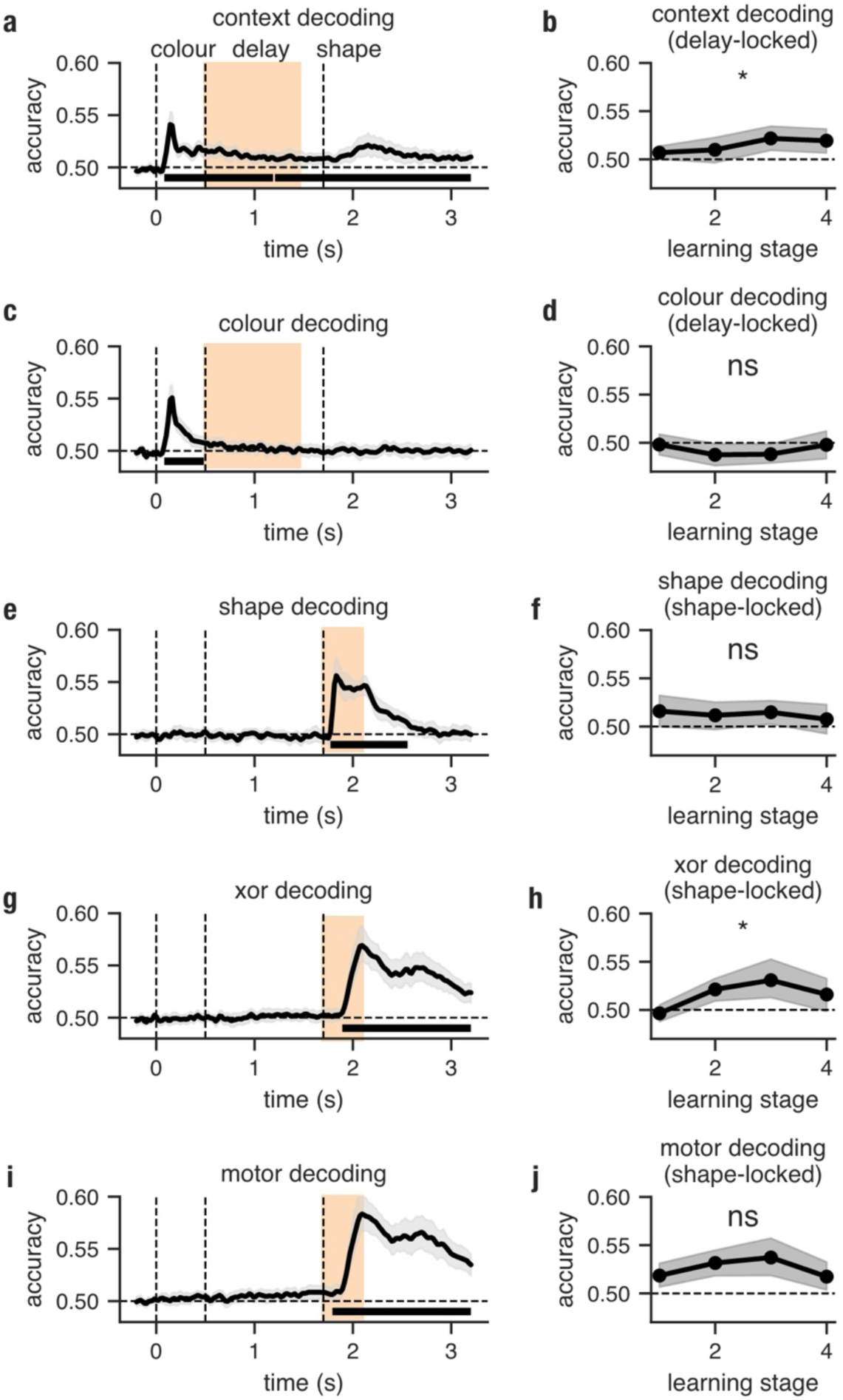
Participants maintain a context signal and construct an XOR representation over learning. **a**, Time-resolved decoding of context; horizontal bars indicate statistical significance; the pale orange area indicates the time windows for which subsequent time-averaged decoding analyses were run (e.g., panel b). Vertical three dashed lines show the onset of the colour, delay, and shape, respectively; **b**, The learning dynamics of the context decoding computed from the time-averaged signal in the delay-locked period. **c-d**, Analogues to a-b but for within-context colour decoding (captures only the physical properties of the colour). **i,j**, **k,l**, and **m,l** Analogues to a-b but for shape, XOR and motor decoding, respectively. The shaded area indicated 95% CIs obtained through random resampling with replacement (*n* = 1000); All p-values for stage 1 vs stage 4 comparisons were calculated using t-tests for dependent samples (***, p < 0.01; **, p <0.01; *, p < 0.05; †, p < 0.1; n.s., not significant).

We hypothesised that working memory processes control the dimensionality of neural representations by selecting features for maintenance. We tested this prediction by exploring the learning dynamics of the colour representation. Importantly, to obtain an isolated measure of colour coding (unrelated to context) we focused on decoding colour that provided the same context information (blue vs pink or green vs khaki). Decoding analyses demonstrated that colour information peaked in the early colour-locked period of the trial and then rapidly declined over time to reach chance levels before the delay-locked period, *cluster* 1: 0.082 − 0.484 *ms*, *p* = 0.006 (**Fig. 2c**). No learning-induced changes to colour decoding were detected between stage 1 and stage 4 in the delay-locked periods, *M* = 0.498 *vs M* = 0.498, *t*(24) = 0.03, *p* = 0.51, *d* = 0.0 (**Fig. 2d**). These results suggest that participants rapidly discarded irrelevant colour information. Only information relevant for performance (context) entered working memory and was maintained.

Time-resolved decoding also revealed that the shape information rapidly peaked, persisted through the presentation period and rapidly fell back to chance after stimulation was over, *cluster* 1: 1.772 − 2.556*s*, *p* = 0.001 (**Fig. 2e**). Furthermore, no learning-induced (stage 1 vs stage 4) changes to shape coding were detected, *M* = 0.516 *vs M* = 0.508, *t*(24) = 0.79, *p* = 0.782, *d* = 0.15 (**Fig. 2f**). XOR and motor decoding followed the same temporal evolution with a later peak (compared to shape) and a slower decay; XOR: *cluster* 1: 1.892 − 3.2 *ms*, *p* = 0.001; motor: *cluster* 1: 1.45 − 1.772 *ms*, *p* = 0.039, *cluster* 2: 1.792 − 3.2 *ms*, *p* = 0.001 (**Fig. 2g** and **Fig. 2i**). In line with our predictions, the XOR signal exhibited a learning-induced increase from stage 1 to stage 4, *M* = 0.496 *vs M* = 0.516, *t*(24) = −1.85, *p* = 0.04, *d* = 0.4 (**Fig. 2h**). No such effect was found for the motor signal (**Fig. 2j**).

### The neural format of the context signal evolves over learning

A defining characteristic of low-dimensional task representations is that they can be easily cross-generalised to different sensory instances of the same task. Guided by this consideration, we next tested whether task features like context, shape, XOR and the motor signal were coded in a similar format across different levels of the irrelevant feature (colour). For example, to explore the cross-colour generalisation of context we trained a classifier on differentiating blue from green and tested it on the discrimination of pink vs khaki, and vice versa (see *Methods, cross-colour generalised decoding* for details). Decoding scores above chance obtained from such an analysis would suggest that the same abstract format of context was used in both sets of colours. It is important to note, however, that the lack of colour coding in the delay- and shape-locked periods indicates that the alignment of discriminant boundaries for each of the task variables (context, shape, XOR and motor coding) across different levels of colour would be trivial in these trial periods.

We found that cross-colour generalised context decoding mirrors the temporal dynamics of simple context decoding only in the shape-locked period, *cluster* 1: 1.892 − 2.315 *ms*, *p* = 0.001, *cluster* 2: 2.335 − 2.516 *ms*, *p* = 0.011 (**Fig. 3a**). There, context transforms into an abstract format only later in the period. Furthermore, the learning-resolved data reveal that contextual information transforms into an abstract cross-generalisable format over learning in the delay-locked period, stage 1 vs stage 4: *M* = 0.502 *vs M* = 0.523, *t*(24) = 1.71, *p* = 0.05, *d* = 0.3 (**Fig. 3b**). Time-resolved cross-colour generalised decoding of shape (*cluster* 1: 1.772 − 2.375 *ms*, *p* = 0.002), XOR (*cluster* 1: 1.892 − 3.2 *ms*, *p* = 0.001) and motor coding (*cluster* 1: 1.289 − 3.2 *ms*, *p* = 0.001; **Fig. 3c,e,g**) and their learning dynamics (**Fig. 3d,f,h**) follow the same pattern of results as observed for simple decoding (**Fig. 2i-n**) with only the XOR exhibiting a learning induced effect, *M* = 0.484 *vs M* = 0.513 (stage 1 vs stage 4), *t*(24) = 2.19, *p* = 0.02, *d* = 0.45. As discussed above, cross-colour generalisation of context in the delay period is already implied by the significant context decoding combined with the absence of irrelevant colour coding noted above. Taken together, these findings imply that participants constructed abstract representations of task features but that the mechanism responsible for this transformation relied heavily on discarding colour information early in trial time. That is, the dimensionality of the task representation was primarily reduced as a function of time in the trial and not changed over learning, through the selection of which information enters working memory. It is worth noting the U-shaped learning dynamic observed across each task variable when learning dynamics were examined (**Fig. 3f,h**). This might be because task-specific geometrical transformations to the neural ctivity might reflect preparation processes and shift with reaction times. Averaging a temporally shifting neural signal using the same time window might thus result in a U-shaped dynamic. We next examined neural representations locked to the response activity to eliminate potential biases from preparation-related shifts.

**Figure 3.**
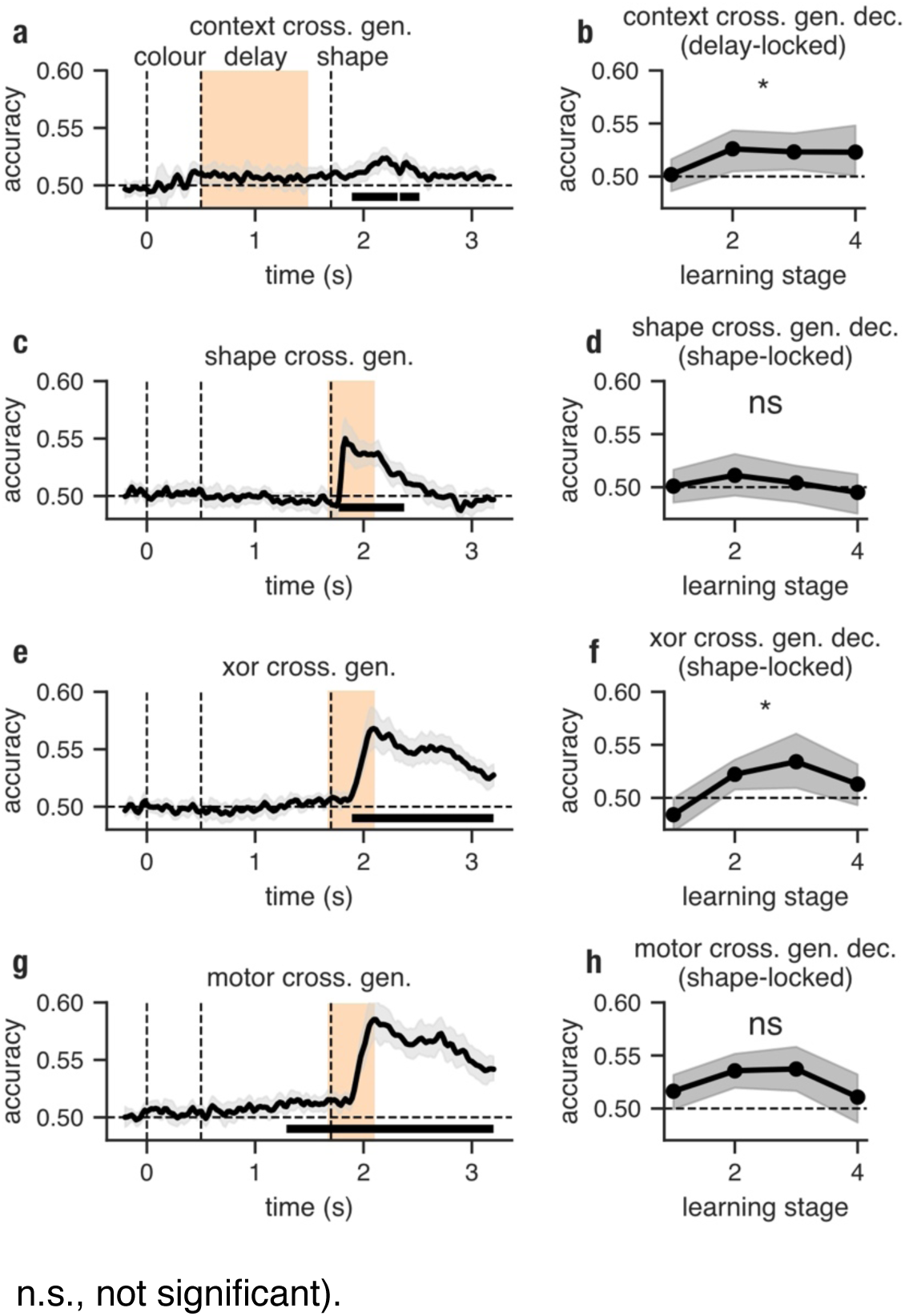
Context and XOR are progressively represented in an abstract format over learning. **a**, Time-resolved cross-colour decoding of context; horizontal bars indicate statistical significance; the pale orange areas indicate the time windows for which subsequent decoding analyses were run (panel b). Vertical three dashed lines show the onset of the colour, delay, and shape, respectively; **b**, The learning dynamics of the cross-colour context decoding computed from the time-averaged signal in the delay period. **c,d** and **e,f**, and **g,h,** Analogues to a-b but for shape, XOR and motor cross-colour decoding, respectively. The shaded area indicated 95% CIs obtained through random resampling with replacement (*n* = 1000); All p-values for stage 1 vs stage 4 comparisons were calculated using t-tests for dependent samples (***, p < 0.01; **, p <0.01; *, p < 0.05; †, p < 0.1; n.s., not significant).

### Analysis of neural geometry reveals evidence of low-dimensional representations at decision time

We hypothesised that the engagement of working memory processes could reflect preparation as part of the necessary computations, and thus lead to a low-dimensional task representation at the decision timepoint. In particular, our previous research showed that non-human primates constructed a low-dimensional and abstract representation of a XOR task over several weeks of training (Wójcik et al., 2022). In that study we found that at the end of learning the task representation was characterised by strong XOR decoding and weak context and shape coding. Furthermore, using cross-generalised decoding we demonstrated that the XOR feature was represented in an abstract format that supported later transfer to a new instance of the same task.

We therefore tested here whether learning in human participants also led to a low-dimensional and abstract task representation at decision time. To test these predictions, we analysed the geometry and dimensionality of averaged neural responses, shortly before the motor response was triggered ([t_-200*ms*_, t_0*ms*_]). We observed that decoding of the XOR feature increases with learning (stage 1 vs stage 4), *M* = 0.516 *vs M* = 0.572, *t*(23) = 4.05, *p* < .001, *d* = 0.77 (**Fig. 4a**). Moreover, the XOR feature became progressively represented in an abstract format over learning as evidenced by the increase in cross-generalised decoding from stage 1 to stage 4, *M* = 0.512 *vs M* = 0.56, *t*(23) = 3.52, *p* = 0.001, *d* = 0.66 (**Fig. 4b**). Additionally, the cross-generalised decoding of context and shape decreased over learning (stage 1 vs stage 4); context: *M* = 0.514 *vs M* = 0.494, *t*(23) = 1.54, *p* = 0.068, *d* = 0.32; shape: *M* = 0.513 *vs M* = 0.493, *t*(23) = 1.86, *p* = 0.038, *d* = 0.41 (**Fig. 4b**). We note that the XOR decoding is colinear with the motor decoding at later stages of learning (i.e., XOR True == Left and XOR False == Right; **Fig. 1c**). However, in early learning (stage 1), when participants have not yet constructed a prominent XOR representation, motor coding is not related to XOR coding. This is corroborated by the above chance-level decoding of motor response in both stage 1 (*M* = 0.592, *t*(23) = 6.65, *p* < .000) and stage 2 (*M* = 0.582, *t*(23) = 6.69, *p* < .000) and no significant change in motor coding over learning (stage 1 vs stage 4), *M* = 0.592 *vs M* = 0.582, *t*(23) = 1.3, *p* = 0.205, *d* = 0.12 (**Fig. 4c**).

**Figure 4.**
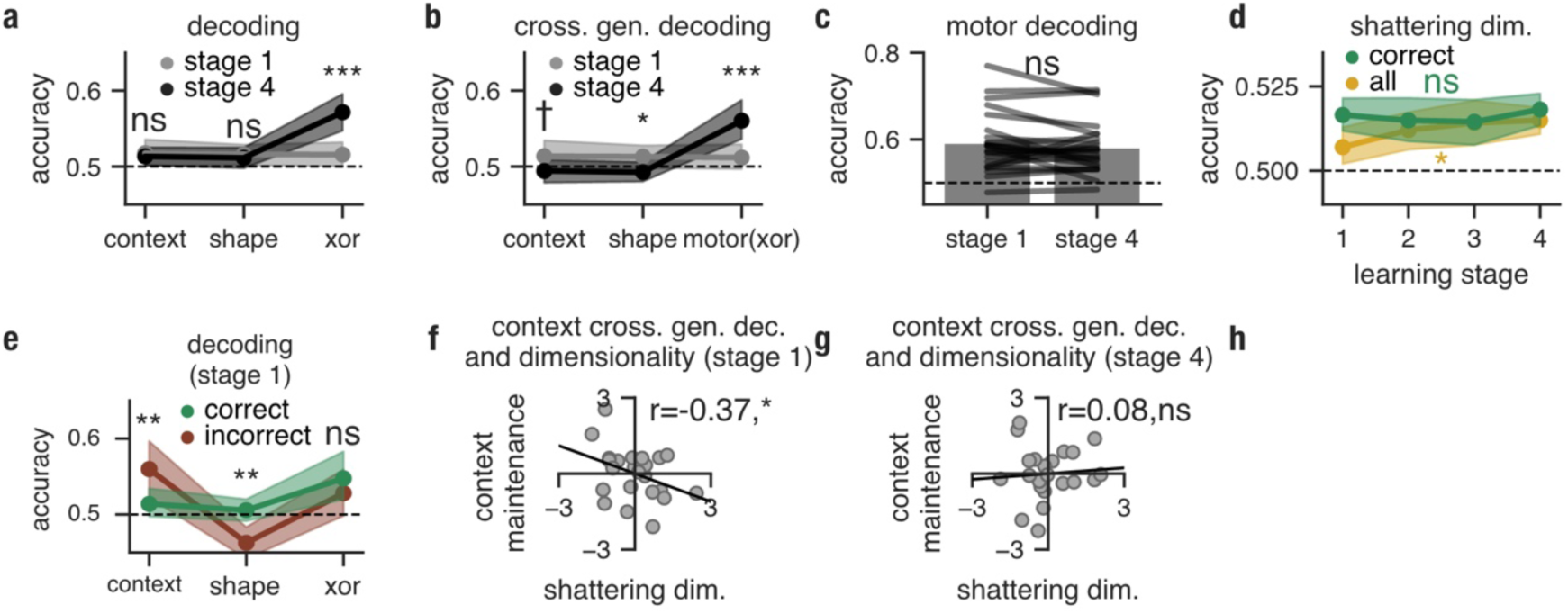
Neural geometry but not dimensionality changes over learning before the decision. **a**, Linear decoding of task variables for learning stages 1 (grey) and 4 (black). **b**, Cross-generalised linear decoding of task variables for learning stages 1 (grey) and 4 (black). **c**, Comparison of motor decoding (response) in stage 1 and stage 4; horizontal black lines represent value pairs for each participant. **d**, The learning dynamic of mean decoding of all possible task dichotomies excluding context, shape and XOR (shattering dimensionality) run on all trials (gold) and only correct trials (green); solid lines indicate the mean and shaded areas 95% confidence intervals over participants, respectively. **e**, Linear decoding of task variables in learning stage 1 run on correct trials (green) and incorrect (red). **f**,**g**, Correlation between normalised mean cross-generalised context decoding scores in the delay-locked period (context maintenance) and the shattering dimensionality computed at the moment of the decision on all trials in stage 1 and stage 4, respectively. All p-values were calculated using a t-test for dependent samples (a-h; l-m) and Pearson’s correlation coefficient (***, *p* < 0.01; **, *p* < 0.01; *, *p* < 0.05; †, *p* < 0.1; *n*. *s*., not significant).

We next explored whether the process of learning influenced the neural dimensionality of the representations used to solve the task. An analysis run on all trials (correct and incorrect combined) indicated that, although shattering dimensionality (see *Method, neural dimensionality* for details) was low throughout the entire experiment, an increase over learning (stage 1 vs stage 4) was observed, *M* = 0.507 *vs M* = 0.515, *t*(22) = 2.73, *p* = 0.012, *d* = 0.5 (gold line, **Fig. 4d**). We hypothesised that low neural dimensionality in learning stage 1 might be related to a high proportion of error trials (Rigotti et al., 2013). That is, a failure in maintaining one of the task features in error trials might have resulted in a low-dimensional representation at the decision timepoint and thus decreased neural dimensionality when all trials were analysed. As the number of error trials was not large enough for a shattering dimensionality analysis, we compared the learning dynamic of shattering dimensionality computed on all trials to the dimensionality dynamic obtained from only correct trials (gold vs green; **Fig. 4d**). This data split revealed that dimensionality in correct trials did not change over learning from stage 1 to stage 4, *M* = 0.518 *vs M* = 0.517, *t*(22) = 0.51, *p* = 0.615, *d* = 0.1. Moreover, a statistically significant difference between shattering dimensionality computed on all trials and computed on correct trials was only detected in learning stage 1, *M* = 0.507 *vs M* = 0.517, *t*(22) = 3.71, *p* = 0.001, *d* = 0.54, i.e., the learning stage containing the most error trials. In summary, these findings suggest that the observed increase in dimensionality reflected the differences between the neural geometry found in correct and incorrect trials rather than being indicative of dimensionality expansion as a learning strategy.

To address this explicitly, we next explored whether incorrect trials in stage 1 are in fact characterised by a weak representation of any of the task features. In line with these predictions, we found that incorrect trials compared to correct trials in stage 1 were characterised by a lower shape decoding score, *M* = 0.46 *vs M* = 0.51, *t*(2182) = 3.02, *p* = 0.004, *d* = 0.73. Interestingly, context decoding was observed to be higher in incorrect trials than in correct trials, *M* = 0.56 *vs M* = 0.51, *t*(18) = 2.64, *p* = .008, *d* = 0.48 (**Fig. 4e**). No significant difference in XOR decoding between incorrect and correct trials was observed, *M* = 0.53 *vs M* = 0.55, *t*(18) = 0.98, *p* = 0.171, *d* = 0.19. These results suggest that the representation used in incorrect trials was dominated by a context representation.

Finally, to test whether working memory engagement resulted in the reduction of neural dimensionality at the moment of the decision, we related the strength of mean cross-generalised context decoding in the delay-locked period to the mean shattering dimensionality (all trials) measured at the moment of the decision at a group level. In line with this hypothesis, we found that the strength of abstract context decoding during maintenance was negatively related to neural dimensionality at the decision time in learning stage 1, *r*(23) = −0.37, *p* = 0.04 (**Fig. 4i**). However, this effect was not present in learning stage 4, *r*(23) = 0.08, *p* = 0.63 (**Fig. 4j**). This suggests that participants that maintained a stronger representation of abstract contextual information during the delay also exhibited lower dimensional representations at decision time.

## Discussion

The recent convergence of neuroscience and machine learning has led to a novel conceptualisation of the role of working memory. It has been proposed that working memory actively participates in improving the efficiency of computation rather than just being a passive data storage system (Dasgupta & Gershman, 2021). Our research echoes this perspective, suggesting that working memory selection shapes neural dimensionality. Notably, these representations closely mirror the energy-efficient neural codes observed in non-human primates following intensive training (Stroud et al., n.d.; Wójcik et al., 2023).

The feature selection mechanism discussed here is based on the assumption that irrelevant features are filtered out after they enter working memory. However, feature selection can also take place at the early perceptual stages through the engagement of selective attention. This is supported by empirical reports demonstrating that the PFC, specifically the inferior temporal junction, can exert top-down control over posterior cortical areas involved in visual processing (Zanto et al., 2010, 2011). Specifically, these frontal areas can enhance or diminish the processing of low-level physical features in visual cortices (from V1 to V5; Ruff et al., 2008) during the early stages of perceptual processing (∼100ms) and thus control what information enters later stages of information processing such as working memory maintenance. As a consequence of attention selection, the diminished representation of the irrelevant feature was observed early in trial time. The early perceptual selection account seems, however, at odds with the described here data. In particular, the strength of colour (irrelevant) and context (relevant) decoding was similar during the presentation period of the coloured cue. Crucially, the irrelevant feature was only discarded during the delay after it entered working memory. Therefore, we believe that the selection mechanism observed here reflects working memory selection rather than selective attention.

In the present study, we also showed that human participants rapidly acquired an abstract representation of context over learning. It should be noted, however, that learning here might not be an effect of slow updates to neural circuitry that accompany building a task representation from scratch. Specifically, it is possible that human participants were already in the possession of a hierarchical XOR representation that could have been used to solve the task and the observed rapid learning speed reflected the alignment of the task representation stored in long-term memory to the new sensory instance. In line with this, our previous research in non-human primates demonstrated that after learning low-dimensional XOR representations, animals were able to rapidly align the already acquired readout dimension to the representation of previously unseen stimuli combinations very early in training (Wójcik et al., n.d.).

While the data presented here shed light on how working memory processes could be utilised to control neural dimensionality, there are potential limitations that should be considered. Firstly, to establish a causal role of working memory selection in reducing the dimensionality of neural activity at the decision timepoint, a control condition could be introduced, whereby the colour information persisted throughout the entire trial. The neural representations and their learning dynamics observed in the current study could be then compared to representation acquired in the second, continuous colour input setting. Secondly, the low difficulty of the task, and consequently a small number of error trials, prevented us from performing a dimensionality analysis on representations that subsequently led to erroneous responses. To address this issue, the difficulty of the task could be increased by reducing the memory delay period. Finally, the motor response signal and the XOR signal were collinear in the study after learning. These two features could be disentangled by introducing a third-level policy and thus a second interaction term. In the current design, the XOR feature represents an interaction between the shape and the colour features. Displaying two abstract stimuli (e.g., “#” and “&” signs) representing the XOR (True vs False) and randomising the sides on which they would be presented (left vs right) would provide a marker of the XOR that is not confounded by motor code.

Building on the results described here, future studies could explore the relationship between working memory processes and low-dimensional task representations using perturbation approaches. Specifically, techniques such as TMS (Rose et al., n.d.) or pinging (Wolff et al., 2017) have been successfully used in the past to reactivate information stored in working memory in an activity-silent format (Stokes, 2015). Using these methods to non-specifically perturb the system right after the stimulus presentation could reveal, for example, whether discarding irrelevant features is related to the decay of neural activity (i.e., in the early delay) (Souza & Oberauer, 2015).

## Methods

### Participants

Twenty-five participants (14 women and 11 men; mean age 23.3 years) were recruited for the study. Before taking part in the experiment, all participants declared their written consent according to the guidelines approved by the MS IDREC of the University of Oxford (approval no R62730/RE001 and R51332). No previous history of neuropsychiatric or neurological disorders was reported by the participants who had either normal or corrected-to-normal vision. All participants received an appropriate monetary reward for their time and effort.

### Procedure and task

Participants were seated in a sound-proof and dimply lit room. Their heads were supported by a chinrest which also ensured a constant viewing distance of 70 *cm*. Visual stimuli were then projected on the screen at a spatial resolution of 1024 *x* 786 pixels with a 60 *Hz* refresh rate. The Psychtoolbox run on MATLAB was used to design and run the task. Each of the conditions (*n* = 8) was repeated 100 times across 20 blocks of 40 trials. The sequence of trials was randomised. Importantly, 5 task variables could have been represented in the task: colour, context, shape, XOR and motor coding (left vs right response).

### Stimuli

The colours of the coloured circles stimulus were chosen using the CIELab colour space (Tkalčič & Tasič, n.d.). To ensure that the colours were approximately isoluminant, the same L parameter was used for every colour. Furthermore, to ensure a circular (equidistant) colour representation, parameters *a* and *b* were fixed with respect to value but not valence. Note that colours were randomly mapped to the XOR rule for every participant before the experiment. This was done to ensure that the initial dissimilarity between colours on a physical level did not influence the learning of the XOR rule at the group level. The shapes were custom-made in Adobe Illustrator.

### EEG acquisition and pre-processing

EEG signal was collected using 61 Ag/AGcl electrodes (EasyCap, Herrshing, Germany) mounted in the cap in line with the 10-20 system (Acharya et al., 2016). The data was amplified and collected at a 1000 *Hz* sampling rate using the NeuroScan SynAmps RT amplifier and the Scan 4.5 software (Compumedics NeuroScan, Charlotte, NC). The electrode system was grounded with an electrode placed at the surface of the right elbow. To allow efficient ocular artefact detection and elimination, horizontal and vertical electrooculography (EOG) was collected. Specifically, two vertical channels were attached to the suborbit and supraorbit of the left eye and two horizontal channels were attached at the exterior canthi of both eyes. The impedance of all channels was checked throughout the duration of the experiment and kept below the level of 10 *kΩ*. After collection, the EEG data were pre-processed using the MNE toolbox that was run on Python 3.5 (Gramfort et al., 2013). Firstly, the data was re-referenced using the right mastoid channel. Then we downsampled the signal to 250*Hz* and used a bandpass filter from 0.01 to 40*Hz*. The signal was subsequently epoched into segments starting −200*ms* and ending 1500*ms*, locked to the onset of colour stimuli and also shape stimuli. For a decision time analysis data was epoched −200 – 0*ms* locked to the button press. Horizontal and vertical eye movement, blinks, ECG and movement artefacts were identified using Independent Component Analysis. Then, we estimated artefact rejection thresholds separately for each participant using the Autoreject algorithm and discarded or repaired corrupt epochs (Jas et al., 2017). We also employed the RANSAC algorithm to identify and repair contaminated sensors (Bigdely-Shamlo et al., 2015). Note that automated pre-processing methods have been shown to be less prone to error than methods based on visual inspection and support transparency, reliability, and scalability of EEG analysis (Jas et al., 2017). Due to technical issues in electrode position digitalisation artifact rejection was based only on automatic threshold estimation in seven participants. Our pre-processing pipeline resulted in a mean value of 728 (*SD* = 85) colour-locked, 728 (*SD* = 83) shape-locked, and 740 (*SD* = 97) response-locked epochs of that were then used in the subsequent analysis phase. For time-averaged analyses these data were divided equally into four learning stages.

### Behavioural analysis

Performance was calculated as the proportion of correct responses in each of the four learning stages. Similarly, reaction times were operationalised as median (over trials) time measured from the onset of the shape stimulus and response button press in each of the four learning stages. Next, we computed three types of switch costs (**Fig. 1e, f, g**) for both performance and reaction times using all trials (**Fig. 1i, j, k** and **m, n, o**, respectively). We first contrasted performance scores and reaction times obtained from *context switch trials* (**Fig. 1e**), where the context on the current trial changed as compared to the previous trial against trials in which the context remained the same (**Fig. 1i, j)**. (Note that here, context denotes the predictive value of the colour, shared between two colours: for example, the square should be mapped to the left button in the context of both the blue and the pink colour; **Fig. 1c**). To test whether context switches impacted performance and reaction times more than shape switches (**Fig. 1f**), we compared context switch trials to *shape switch trials* (**Fig. 1f, k)**. Finally, context switch trials were compared to *colour switch trials* (**Fig. 1g**), where the colour on the current trial changed as compared to the previous trials but the context remained the same (**Fig. 1g, l**). This was done to measure a context-independent impact of a sensory switch on performance and reaction times. Importantly, as frequentist statistics were used in the behavioural analysis, reaction times distribution were transformed using a log transform to approximate a normal distribution.

### Decoding

To determine the amount of information represented for each task variable by the brain linear discriminant analysis (LDA) decoders were employed. The inherent presence of noise and spatial correlations observed in the EEG signal can make decoding particularly difficult. Specifically, because electrical potentials can propagate freely through brain tissue, the same source signal can be measured by multiple EEG sensors leading to a spatial covariance structure that is prevalent in the observed signal. Additionally, the process of referencing introduces artefacts present at the reference site to every sensor in the dataset. To account for these characteristics a LDA decoder with shrinkage was used (for similar shrinkage applications in decoding see Wolff et al., 2017). The shrinkage factor was chosen based on the Ledoit-Wolf lemma (Ledoit & Wolf, 2004). The decoding analysis was run within each participant for each time point separately. First, trials were split randomly into equal test and train sets. Next, an LDA classifier was trained on the train data and subsequently tested on the test data (and vice-versa). The scores obtained from these two procedures were then averaged. This was repeated for 10 random 50%/50% splits and the scores were again averaged. Importantly, to obtain an estimate of colour decoding that was not biased by context information (i.e., an estimate of pure irrelevant information) we run two separate decoders that differentiated colours within the same context: (1) colour 1 vs colour 3 (context 1), and (2) colour 2 from colour 4 (context 1). The scores were then averaged. Decoding analyses were also run on time average data. For this purpose, learning resolved data (4 learning stages) were time-averaged (trial time) in the colour-locked period ([t_100*ms*_, t_500*ms*_], colour-locked), delay-locked period ([t_500*ms*_, t_1500*ms*_], colour-locked), and shape-locked period ([t_100*ms*_, t_500*ms*_], shape-locked). When analysing neural activity at the moment of decision the EEG signal was averaged in the response-locked period ([t-_200*ms*_, t_0*ms*_]).

### Neural dimensionality

Neural dimensionality was operationalised in this study as shattering dimensionality, a metric that has been applied in several previous experiments (Bernardi et al., 2020; Rigotti et al., 2013; Wójcik et al., n.d.). We utilised the same decoding method mentioned earlier but averaged the results across all 35 potential dichotomies possible given the task’s structure. This task involves three linear variables: colour, context, and shape. When combined, they create a cube in the input space, which can be divided in 35 unique ways, each representing two sets of 4 vertices. We determined how decodable each dichotomy was and observed the average decoding accuracy over time.

### Full cross-generalised decoding

To explore the neural geometry of the task information represented by the participants at decision time we used cross-generalised LDA decoding (Bernardi et al., 2020; Wójcik et al., 2023). Similarly as previous studies, trials were split according to the to-be-decoded target variable labels as well as the splitting variable labels. That is, the splitting variable provided the training and test instances of the data for the decoding of the target variable. The task examined here had three linear input variables (colour, context and shape) which resulted in 36 possible cross. gen axes per each of the task variables. That is, the labels for 8 conditions (2 *colours x* 2 *shapes x* 2 *widt*h*s*) are split into two sets of four labels (by colour, shape, or width). To identify training and testing exemplars within each of the sets, every possible binary combination of condition labels was determined (4 *c*h*oose* 2). This yielded 6 combinations for each of the sets. Thirty-six unique train-test splits (cross-generalisation axes) can be achieved by combining these two sets. Decoding accuracy scores estimated by running linear LDA decoders for each of these splits were then averaged to obtain a general cross-generalisation score of the analysed target variable.

### Cross-colour generalised decoding

The examine whether the task-irrelevant feature (colour) was discarded by the cortex and therefore influenced neural geometry a simpler format of cross-generalised LDA decoding was employed. Specifically, we tested whether learning the task resulted in representations that were shared across different levels of the irrelevant feature (colour). This resulted in four possible binary decoding problems (e.g., when performing cross-generalised decoding for the context variable we can: (1) train on differentiating colour 1 (context 1) from colour 2 (context 2) and test on differentiating colour 3 (context 1) from colour 4 (context 2), (2) train on differentiating colour 3 (context 1) from colour 4 (context 2) and test on differentiating colour 1 (context 1) from colour 2 (context 2), (3) train on differentiating colour 1 (context 1) from colour 4 (context 2) and test on differentiating colour 2 (context 2) from colour 3 (context 1), and (4) train on differentiating colour 2 (context 2) from colour 3 (context 1) and test on differentiating colour 1 (context 1) from colour 4 (context 2); these four decoding scores were then averaged). Using this procedure, we explored the cross-colour generalisation potential of the context, shape, XOR and motor coding variables.

### Significance testing

Time-resolved one-sample t-tests (one-tailed) were used to test for statistical significance of the temporally resolved decoding accuracy scores. To correct for family-wise errors a cluster-based correction was employed (Maris & Oostenveld, 2007). To test for learning-induced changes in accuracy, reaction times, decoding and selectivity measures we compared data from learning stage 1 to data from learning stage 4 using a t-test for dependent samples. Furthermore, shaded areas in all figures represent 95% confidence intervals obtained through a resampling with replacement procedure (*n* = 1000). In analyses where decoding scores were compared to the chance level, one-sample t tests were used. Test statistics were supplemented with Cohen’s *d* effect size estimation. Two participants were excluded from the response-locked analyses due to trigger failure and an insufficient number of error trials.

